# Sex differences in lung imaging and SARS-CoV-2 antibody responses in a COVID-19 golden Syrian hamster model

**DOI:** 10.1101/2021.04.02.438292

**Authors:** Santosh Dhakal, Camilo A. Ruiz-Bedoya, Ruifeng Zhou, Patrick S. Creisher, Jason S. Villano, Kirsten Littlefield, Jennie Ruelas Castillo, Paula Marinho, Anne Jedlicka, Alvaro A. Ordonez, Natalia Majewski, Michael J. Betenbaugh, Kelly Flavahan, Alice L. Mueller, Monika M. Looney, Darla Quijada, Filipa Mota, Sarah E. Beck, Jacqueline Brockhurst, Alicia Braxton, Natalie Castell, Franco R. D’Alessio, Kelly A. Metcalf Pate, Petros C. Karakousis, Joseph L. Mankowski, Andrew Pekosz, Sanjay K. Jain, Sabra L. Klein, for the Johns Hopkins COVID-19 Hamster Study Group

## Abstract

In the ongoing coronavirus disease 2019 (COVID-19) pandemic caused by the severe acute respiratory syndrome coronavirus 2 (SARS-CoV-2), more severe outcomes are reported in males compared with females, including hospitalizations and deaths. Animal models can provide an opportunity to mechanistically interrogate causes of sex differences in the pathogenesis of SARS-CoV-2. Adult male and female golden Syrian hamsters (8-10 weeks of age) were inoculated intranasally with 10^5^ TCID_50_ of SARS-CoV-2/USA-WA1/2020 and euthanized at several time points during the acute (i.e., virus actively replicating) and recovery (i.e., after the infectious virus has been cleared) phases of infection. There was no mortality, but infected male hamsters experienced greater morbidity, losing a greater percentage of body mass, developing more extensive pneumonia as noted on chest computed tomography, and recovering more slowly than females. Treatment of male hamsters with estradiol did not alter pulmonary damage. Virus titers in respiratory tissues, including nasal turbinates, trachea, and lungs, and pulmonary cytokine concentrations, including IFNβ and TNFα, were comparable between the sexes. However, during the recovery phase of infection, females mounted two-fold greater IgM, IgG, and IgA responses against the receptor-binding domain of the spike protein (S-RBD) in both plasma and respiratory tissues. Female hamsters also had significantly greater IgG antibodies against whole inactivated SARS-CoV-2 and mutant S-RBDs, as well as virus neutralizing antibodies in plasma. The development of an animal model to study COVID-19 sex differences will allow for a greater mechanistic understanding of the SARS-CoV-2 associated sex differences seen in the human population.

**Importance:** Men experience more severe outcomes from COVID-19 than women. Golden Syrian hamsters were used to explore sex differences in the pathogenesis of a human clinical isolate of SARS-CoV-2. After inoculation, male hamsters experienced greater sickness, developed more severe lung pathology, and recovered more slowly than females. Sex differences in disease could not be reversed by estradiol treatment in males and were not explained by either virus replication kinetics or the concentrations of inflammatory cytokines in the lungs. During the recovery period, antiviral antibody responses in the respiratory tract and plasma, including to newly emerging SARS-CoV-2 variants, were greater in females than male hamsters. Greater lung pathology during the acute phase combined with reduced antiviral antibody responses during the recovery phase of infection in males than females illustrate the utility of golden Syrian hamsters as a model to explore sex differences in the pathogenesis of SARS-CoV-2 and vaccine-induced immunity and protection.

**One Sentence Summary:** Following SARS-CoV-2 infection, male hamsters experience worse clinical disease and have lower antiviral antibody responses than females.

## Introduction

At the start of the coronavirus disease 2019 (COVID-19) pandemic, early publications from Wuhan, China (1, 2) and European countries (3) began reporting male biases in hospitalization, intensive care unit (ICU) admissions, and mortality rates. Ongoing real-time surveillance (4) and meta-analyses of over 3 million cases of COVID-19 (5) continue to show that while the incidence of COVID-19 cases are similar between the sexes, adult males are almost 3-times more likely to be admitted into ICUs and twice as likely to die as females. Differential exposure to the severe acute respiratory syndrome coronavirus 2 (SARS-CoV-2) is likely associated with behaviors, occupations, comorbidities and societal and cultural norms (i.e., gender differences) that impact the probability of exposure, access to testing, utilization of healthcare, and risk of disease (6-8). This is distinct but also complementary to biological sex differences (i.e., sex chromosome complement, reproductive tissues, and sex steroid hormone concentrations) that can also impact susceptibility and outcomes from COVID-19 (9, 10). While exposure to SARS-CoV-2 may differ based on gender, the increased mortality rate among males in diverse countries and at diverse ages likely reflect biological sex. Studies have shown that in males, mutations in X-linked genes (e.g., *TLR7*) resulting in reduced interferon signaling (11), elevated proinflammatory cytokine production (e.g., IL-6 and CRP) (2, 12), reduced CD8+ T cell activity (e.g., IFN-γ) (13), and greater antibody responses (i.e., anti-SARS-CoV-2 antigen-specific IgM, IgG, and IgA, and neutralizing antibodies) (14) are associated with more severe COVID-19 outcomes as compared with females. Because COVID-19 outcomes can be impacted by both gender and biological sex, consideration of the intersection of these contributors is necessary in human studies (15).

Animal models can mechanistically explore sex differences in the pathogenesis of SARS-CoV-2 independent of confounding gender-associated factors that impact exposure, testing, and use of healthcare globally. Transgenic mice expressing human ACE2 (K18-hACE2) are susceptible to SARS-CoV-2 and in this model, males experience greater morbidity than females, despite having similar viral loads in respiratory tissues (e.g., nasal turbinates, trachea, and lungs) (16, 17). Transcriptional analyses of lung tissue revealed that inflammatory cytokine and chemokine gene expression is greater in males than females early during infection, and these transcriptional patterns show a stronger correlation with disease outcomes among males than females (16, 17). In addition to utilizing hACE2 mice, mouse-adapted strains of SARS-CoV-2 have been developed and can productively infect wild-type mice but have not yet been used to evaluate sex-specific differences in the pathogenesis of disease (18-20).

Golden Syrian hamsters are also being used as an animal model of SARS-CoV-2 pathogenesis because they are susceptible to human clinical strains of viruses, without the need for genetic modifications in either the host or virus. While studies have included males and females in analyses of age-associated differences in the pathogenesis of SARS-CoV-2 (21), few studies have specifically evaluated males vs. females to better understand sex differences in disease. There are studies of golden Syrian hamsters that have included male and female hamsters but did not have sufficient numbers of animals to accurately compare the sexes (22). Sex differences are not reported in either viral RNA, infectious virus, or cytokine mRNA expression at a single time point (i.e., 4 days post-infection) in the lungs of golden Syrian hamsters (23). There is a gap in the literature of studies designed to rigorously test the hypothesis that biological sex alters disease severity and immune responses after SARS-CoV-2 infection.

## Results

### Males experience greater morbidity than females following SARS-CoV-2 infection, which cannot be reversed by estradiol (E2) treatment

Intranasal inoculation of human clinical isolates of SARS-CoV-2 causes productive infection in golden Syrian hamsters (24-26). To test the hypothesis that SARS-CoV-2 infection results in sex differences in disease outcomes, adult male and female golden Syrian hamsters were infected with 10^5^ TCID_50_ of virus and changes in body mass were monitored for 28 days post-inoculation (dpi). Mortality was not observed in either sex, but infected hamsters progressively lost body mass during the first week before starting to recover (**Figure 1A**). The peak body mass loss in female hamsters was observed at 6 dpi (−12.3±1.8%), whereas peak mass loss in male hamsters was observed at 7 dpi (−17.3±1.9%). The percentage of body mass loss was significantly greater in male than female hamsters at 8 to 10 dpi and throughout the recovery period (*p*<0.05; **Figure 1A**). Recovery to baseline body mass after SARS-CoV-2 infection occurred within 2 weeks for female and at 3 weeks for male hamsters (**Figure 1A**).

**Figure 1:**
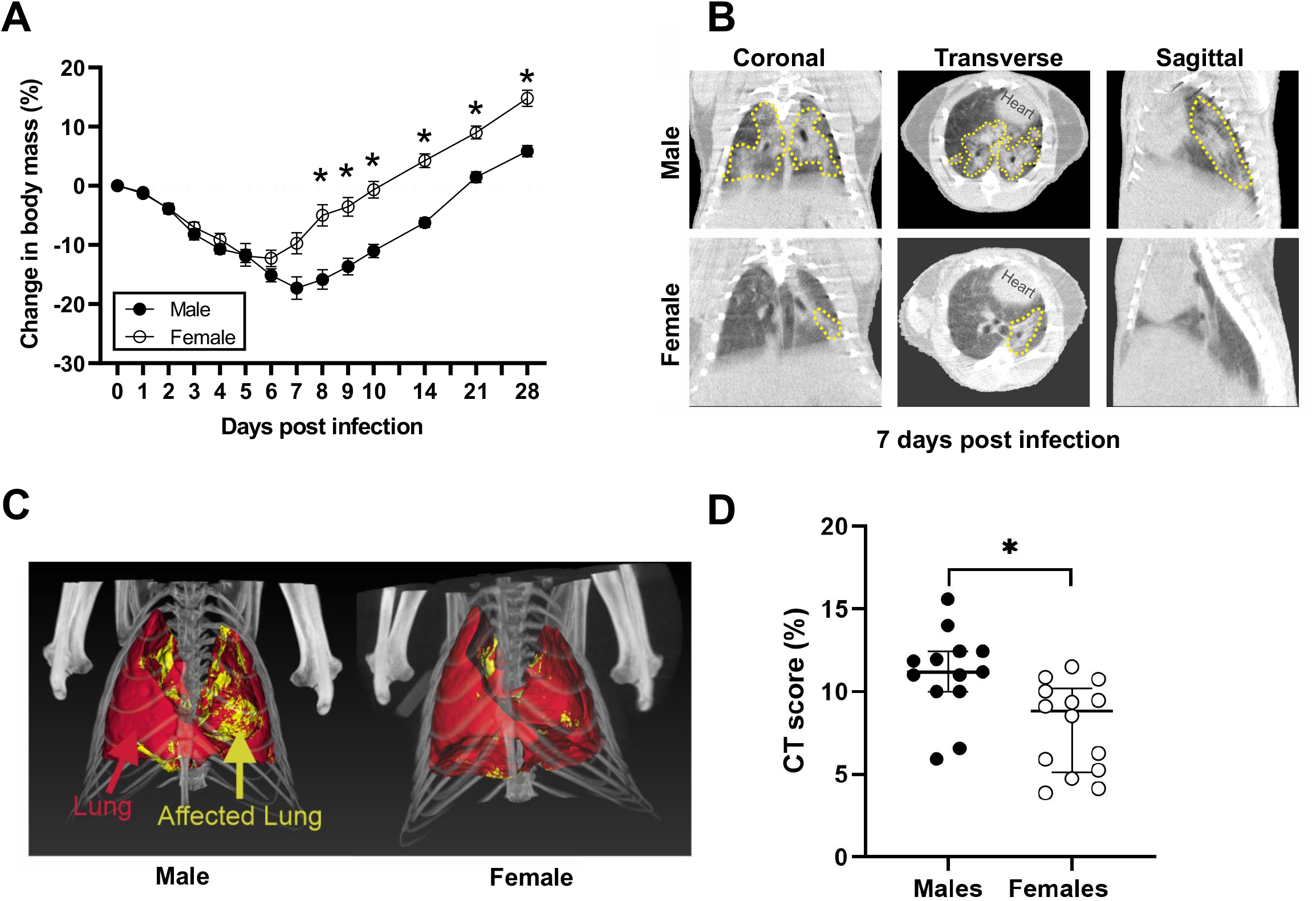
SARS-CoV-2 infected male hamsters experience greater disease than females. To evaluate morbidity, the percent change in body mass from pre-inoculation was measured up to 28 dpi (A). Representative coronal, transverse, and sagittal chest CT from SARS-CoV-2-infected male and female animals are shown (B). Lung lesions (GGO, consolidation and air bronchogram) are marked by the dashed yellow lines. Maximum intensity projections (MIP) marking total (red) and diseased lung (yellow) for both males and females are shown (C). The CT score is higher in male versus female hamsters at 7 dpi (D). Weights are represented as mean ± standard error of the mean from two independent replications (n = 9-10/group), and significant differences between groups are denoted by asterisks (*p<0.05) based on two-way repeated measures ANOVA followed by Bonferroni’s multiple comparison (A). Chest CT data is represented as median ± interquartile range from two independent replication (13-14/group) and significant differences between groups are denoted in asterisk (*p<0.05) based on unpaired two-tailed Mann-Whitney test (D).

To evaluate pulmonary disease in SARS-CoV-2-infected males and females, chest computed tomography (CT) was performed at the peak of lung disease (7 dpi). As previously reported by others (26), multiple and bilateral mixed ground-glass opacities (GGO) and consolidations were detected in both females and males (**Figure 1B** and **Supplementary Figure 1**). In order to reduce bias in the visual assessment, we developed an unbiased approach to quantify lung disease by chest CT. Volumes of interest (VOIs) were drawn to capture total and diseased (pneumonic) lung volumes (**Figure 1C**). As reported in COVID-19 patients who underwent CT (27, 28), there was significantly more disease in the lung of male versus female hamsters (*p*<0.05) (**Figure 1D**). These results indicate that infected male hamsters developed more severe disease, including more extensive lung injury, than females.

Previous studies show that estrogens, including but not limited to estradiol (E2), are anti-inflammatory and can reduce pulmonary tissue damage following respiratory infections, including with influenza A viruses or *Streptococcus pneumoniae* (29-31). To test the hypothesis that E2 could dampen inflammation and pulmonary tissue damage to improve outcomes in male hamsters, males received either exogenous E2 capsules or placebo capsules prior to SARS-CoV-2 infection. Plasma concentrations of E2 were significantly elevated in E2-treated males compared with placebo-treated males (*p*<0.05; **Figure 2A**) and were well within the normal range of plasma concentrations of E2 in cyclic female hamsters (30-700pg/mL) (32). Animals were followed for 7 dpi and changes in body mass and chest CT score were quantified. There was no effect of E2-treatment on morbidity as placebo- and E2-treated males had equivalent percentages of body mass loss (**Figure 2B**). CT findings noted in E2-treated males were similar to those noted in placebo-treated males (**Supplementary Figure 1**) and chest CT scans revealed in CT score between groups (**Figure 2C**). Moreover, histopathology demonstrated similar cell infiltration and pneumonic areas between groups (**Figure 2D**). From these data, we conclude that the treatment of gonadally-intact males with E2 did not improve morbidity or pulmonary outcomes from SARS-CoV-2 infection.

**Figure 2:**
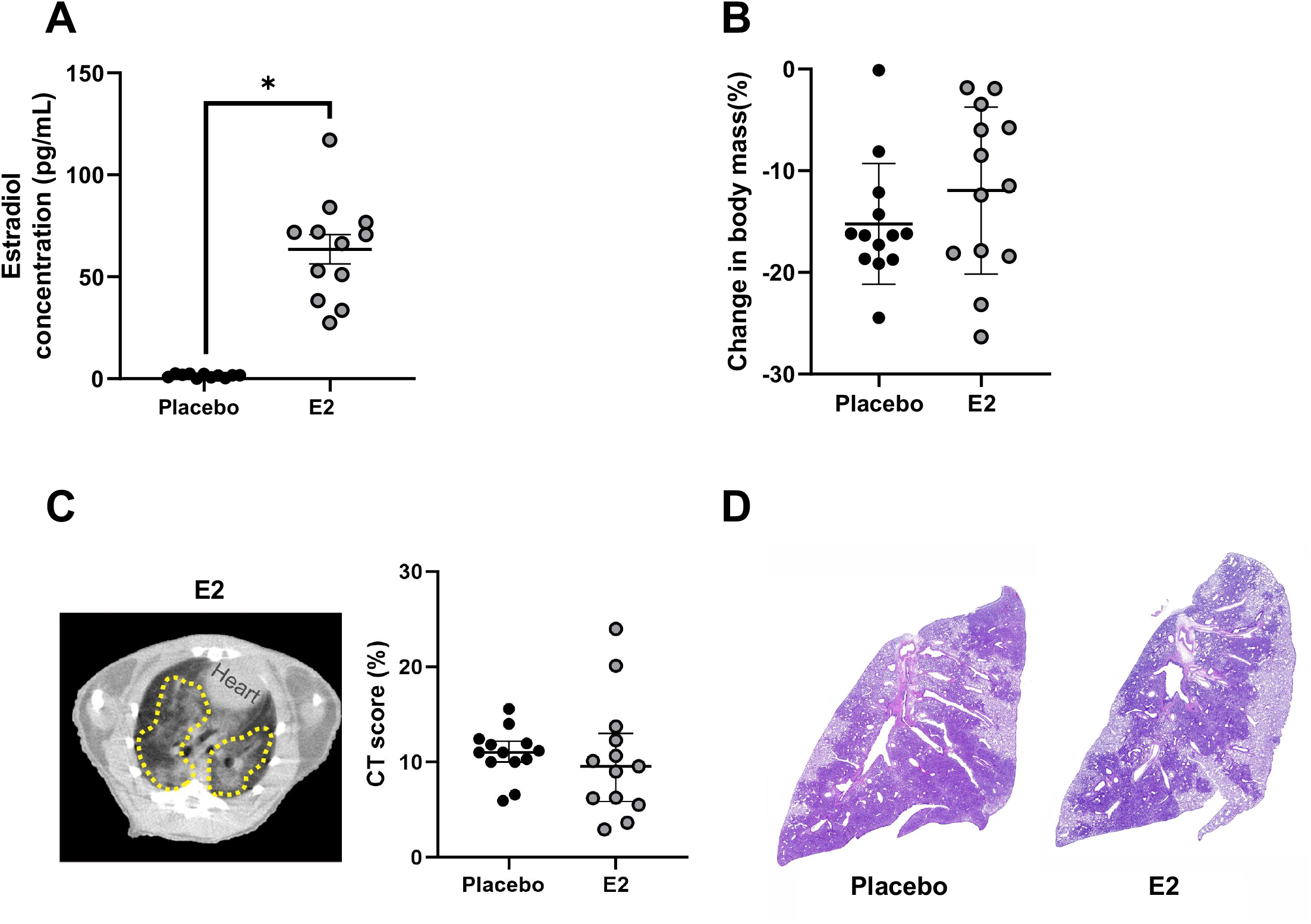
SARS-CoV-2 infected male hamsters treated with estradiol (E2) developed similar lung pathology as placebo-treated males. Male hamsters were treated with E2 capsules or placebo capsules prior to SARS-CoV-2 infection. Estrogen levels were quantified in plasma at 7 dpi (A). Change in body mass for E2-and placebo-treated males were quantified (B). CT score shows no difference between E2-treated males and placebo-treated males (C). Histopathology (H&E) in a representative SARS-CoV-2-infected placebo-treated male and E2-treated male hamster lungs at 4X magnification are shown (D). The dashed yellow lines indicate lung lesions (GGO, consolidations and air bronchogram). E2 concentrations represented as mean ± standard error of the mean of two independent experiments (n=11-12/group) and significant differences between groups are denoted in asterisk (*p<0.05) based on two-tailed unpaired t-test (A). Weight represented as mean ± standard error of the mean of two independent experiments (n=13/group) (B). Chest CT data represented as median ± interquartile range (IQ) from two independent experiments (n = 13/group) (C).

### SARS-CoV-2 replication kinetics are similar between the sexes

To test the hypothesis that male-biased disease outcomes were caused by increased virus load or faster replication kinetics, subsets of infected male and female hamsters were euthanized at 2, 4, or 7 dpi and infectious virus titers were measured in the respiratory tissue homogenates. The peak infectious virus titers in the nasal turbinates (**Figure 3A**), trachea (**Figure 3B**), and lungs (**Figure 3C**) were detected at 2 dpi, decreased at 4 dpi, and was cleared at 7 dpi. There were no sex differences in either peak virus titers or clearance of SARS-CoV-2 from any of the respiratory tissues tested (**Figure 3A-C**). Although the infectious virus was cleared from the respiratory tract of most of the hamsters by 7 dpi (**Figure 3A-C**), viral RNA was still detectable in the lungs at 14 dpi in all of the SARS-CoV-2 infected hamsters, with no differences between the sexes (**Figure 3D**). These data illustrate that sex differences in the disease phenotype are not due to differences in infectious virus loads or persistence of viral RNA.

**Figure 3:**
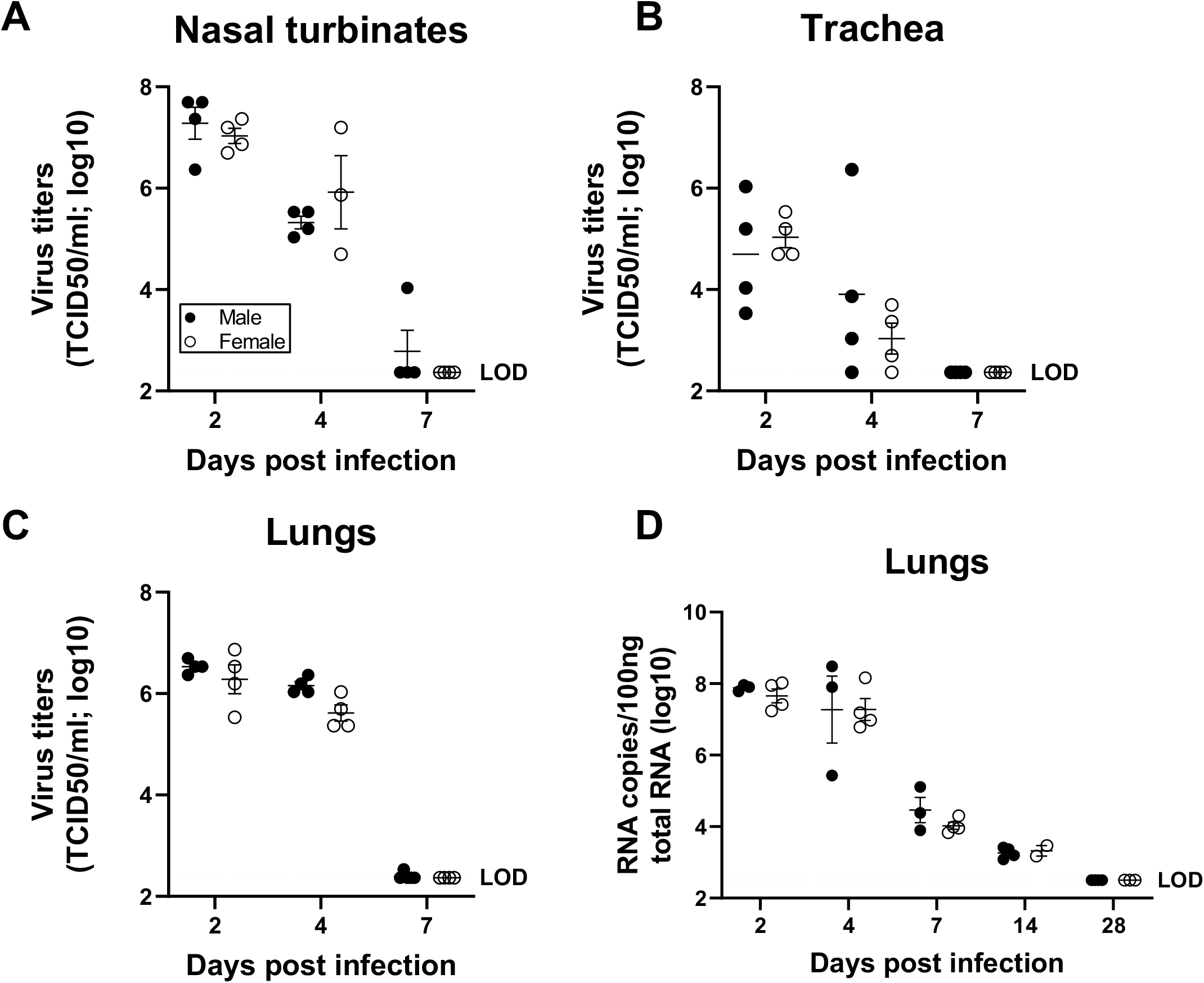
Virus titers were comparable in the respiratory system of SARS-CoV-2 infected male and female hamsters. Adult (8-10 weeks) male and female golden Syrian hamsters were infected with 10^5^ TCID_50_ of SARS-CoV-2. Infectious virus titers in the homogenates of nasal turbinates (A), trachea (B), and lungs (C), were determined by TCID_50_ assay on 2, 4, and 7 dpi. Likewise, virus RNA copies in 100ng of total RNA were tested in the lungs of infected hamsters at 2, 4, 7, 14 and 28 dpi (D). Data represent mean ± standard error of the mean from one or two experiment(s) (n = 3-5/group) and were analyzed by two-way ANOVA (mixed-effects analysis) followed by Bonferroni’s multiple comparison test.

### Cytokine concentrations in the lungs are comparable between the sexes

To test whether local or systemic cytokine activity differed between the sexes, concentrations of cytokines were measured in lung and spleen homogenates at 2, 4, or 7 dpi. Sex differences were not observed 2-7 dpi in the concentrations of IL-1β, TNF-α,, IL-6, IFN-α, IFN-β, IFN-γ, or IL-10 in either lung (**Figure 4A-F**) or spleen (**Supplementary Table 1**) homogenates. In both male and female hamsters, lung concentrations of IL-1β (**Supplementary Figure 2A**), TNF-α (**Supplementary Figure 2B**), IFN-α (**Supplementary Figure 2D**), and IFN-β (**Supplementary Figure 2E**), but not IL-6 (**Supplementary Figure 2C**), IFN-γ (**Supplementary Figure 2F**), or IL-10 (**Supplementary Table 1)**, were greater in samples from infected as compared with sex-matched mock-infected hamsters (*p*<0.05 in each case). In contrast, there was no effect of infection on the concentration of cytokines in the spleen (**Supplementary Table 1**). To determine if cytokine concentrations correlated with virus titers from the same lung homogenates, Spearman correlational analyses were performed and revealed that concentrations of TNFα were positively associated with virus titers at 2 dpi ((*p*<0.05; **Supplementary Figure 3B**) and concentrations of IFNβ were negatively associated with virus titers at 4 dpi (*p*<0.05; Supplementary **Figure 4E**). The concentrations of other cytokines measured at either 2 or 4 dpi were not associated with virus titers in the lungs (**Supplementary Figures 3-4**) Taken together, these data provide no evidence that male-biased disease outcomes are caused by differential production of cytokines in response to SARS-CoV-2 during acute infection.

**Figure 4:**
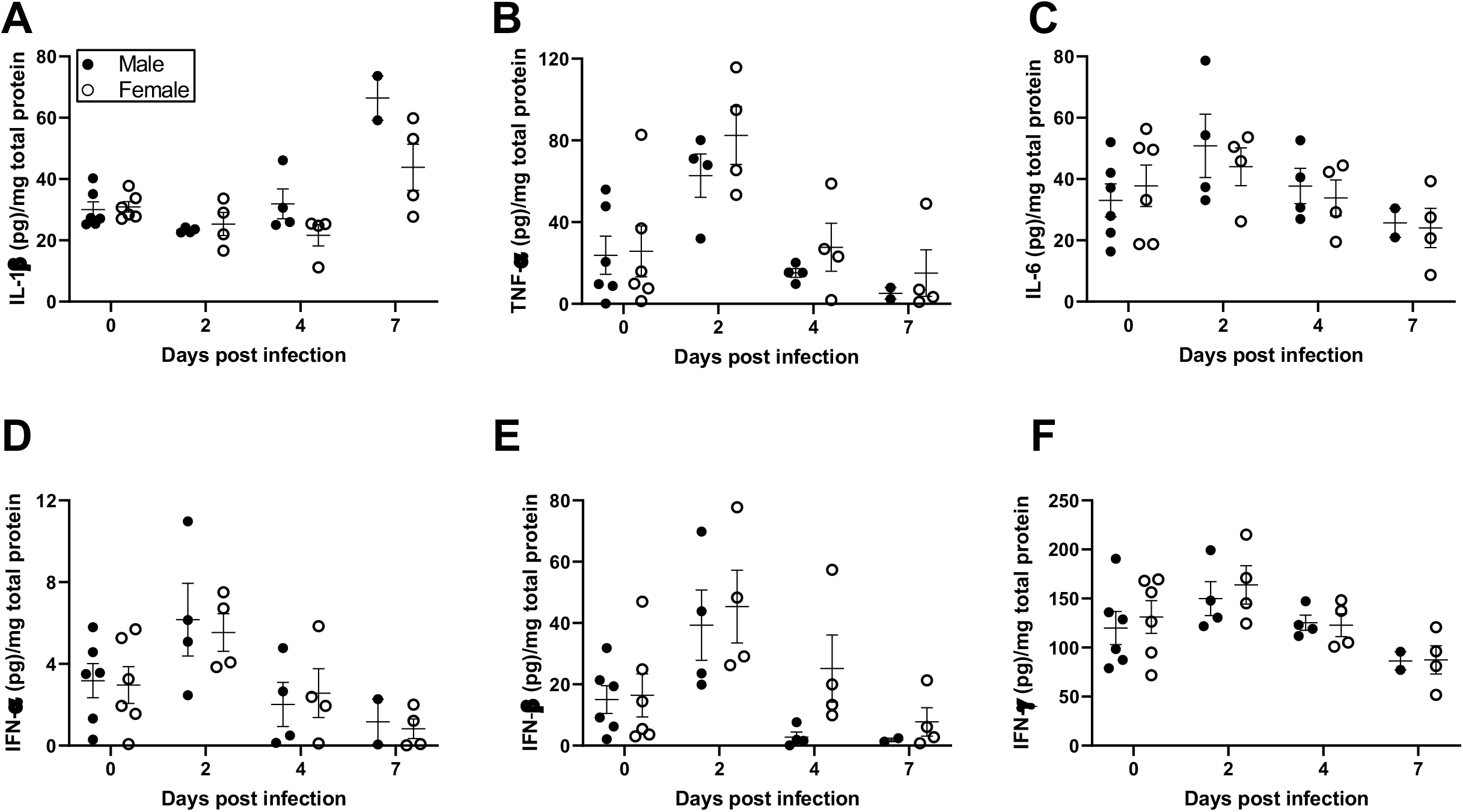
Cytokine responses in the lungs of SARS-CoV-2 infected male and female hamsters were comparable. Adult (8-10 weeks) male and female golden Syrian hamsters were infected with 10^5^ TCID_50_ of SARS-CoV-2. Subsets of animals were euthanized at different dpi and IL-1β (A), TNF-α (B), IL-6 (C), IFN-α (D), IFN-β (E), and IFN-γ (F) cytokine concentrations (pg/mg total protein) were determined in the lungs by ELISA. Mock-infected animal samples from different dpi were presented together as 0 dpi. Data represent mean ± standard error of the mean from one or two independent experiments (n = 2-6/group/sex) and were analyzed by two-way ANOVA (mixed-effects analysis) followed by Bonferroni’s multiple comparison test.

### Female hamsters develop greater antibody responses than males during SARS-CoV-2 infection

To evaluate whether females developed greater antiviral antibody responses than males, as is observed in response to influenza A viruses (33), we measured virus-specific immunoglobulins as well as neutralizing antibody (nAb) titers in plasma and respiratory samples collected throughout the course of infection. To begin our evaluation, we inactivated SARS-CoV-2 virions to analyze plasma IgG that recognize diverse virus antigens. Anti-SARS-CoV-2 IgG titers were detected within a week post-infection, with females developing greater antibody titers than males at 21 and 28 dpi (*p*<0.05; **Figure 5A**). Using live SARS-CoV-2, we measured nAb titers in plasma, which were detectable 7-28 dpi, with females having or trending towards significantly greater titers than males at 14-28 dpi (*p*<0.05; **Figure 5B**).

**Figure 5:**
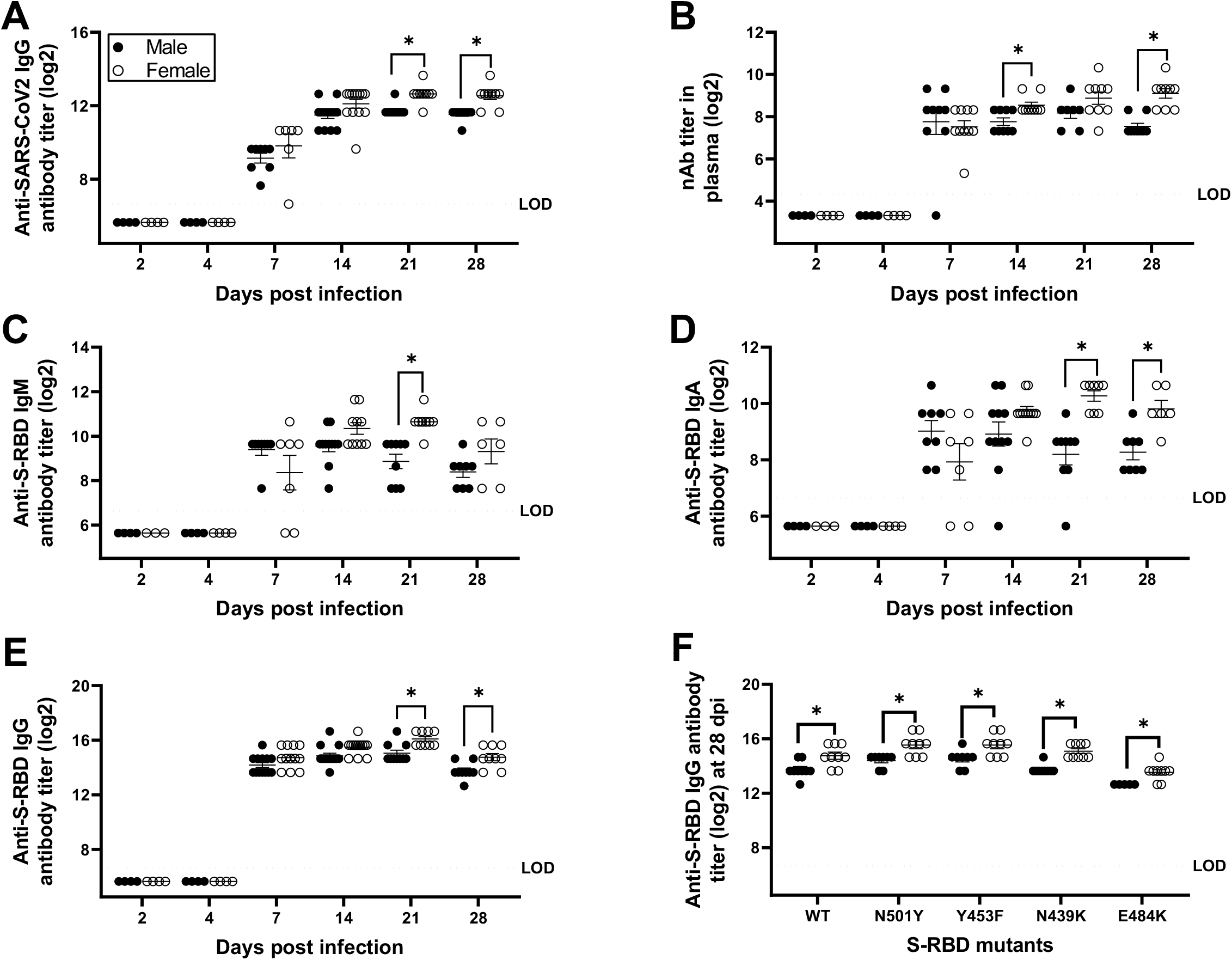
Antibody responses in the plasma of SARS-CoV-2 infected female hamsters were greater than males. Plasma samples were collected at different dpi and IgG antibody responses against whole inactivated SARS-CoV-2 virions (A); virus neutralizing antibody responses (B); and S-RBD-specific IgM (C), IgA (D), and IgG (E) antibodies were determined. Likewise, cross-reactive IgG antibodies against mutant S-RBDs (viz. N501Y, Y453F, N439K, and E484K) were evaluated in plasma at 28 dpi (F). Considering similar antibody responses at 6 and 7 dpi, values were presented together as 7 dpi. Data represent mean ± standard error of the mean from two independent experiments (n = 4-14/group/sex) and significant differences between groups are denoted by asterisks (*p<0.05) based on two-way ANOVA (mixed-effects analysis) followed by Bonferroni’s multiple comparison test.

SARS-CoV-2 infection induces robust antibody responses against the spike or receptor-binding domain of the spike protein (S-RBD) in humans and in animal models (14, 34, 35). S-RBD-specific IgM (**Figure 5C**), IgA (**Figure 5D**), and IgG (**Figure 5E**) antibodies were detected in plasma within a week post-infection. In plasma, anti-S-RBD IgM antibody titers were significantly greater in females than males at 21 dpi (*p*<0.05; **Figure 5C**), and anti-S-RBD IgA and IgG antibody titers were significantly greater in females than males at 21 and 28 dpi (*p*<0.05; **Figure 5D-E**). Variants of SARS-CoV-2 due to mutations in the RBD of spike protein, including the N501Y variant, were first reported in the United Kingdom and subsequently circulated worldwide (36). The mink variant (Y453F), European variant (N439K), and South African/Brazilian variants (E484K) have raised concerns over increased transmissibility and escape from host immune responses (37, 38). Considering the emergence of novel variants, we tested the hypothesis that females would have greater cross-reactive antibody responses to SARS-CoV-2 variants. Similar to wild type S-RDB (**Figure 5E**), IgG antibody titers against the S-RBD mutants N501Y, Y453F, N439K, and E484K were significantly greater in female than male hamsters (*p*<0.05 in each case; **Figure 5F**). Overall, IgG responses to the E484K, but not the N501Y variant, were significantly lower in both sexes as compared with responses to the wild-type S-RBD (*p*<0.05 for main effect of variant; **Figure 5F**).

Local antibody responses at the site of infection are critical for SARS-CoV-2 control and recovery (39, 40). Anti-S-RBD-IgM titers were greatest in the lungs at 7 dpi, being significantly greater in female than male hamsters (*p*<0.05; **Figure 6A**). A cornerstone of mucosal humoral immunity is IgA and anti-S-RBD IgA titers peaked at 7 dpi, with a trend for higher titers in females than males (*p*=0.07; **Figure 6B**). By 28 dpi, females still had detectable anti-S-RBD IgA titers in their lungs, whereas males did not (*p*<0.05; **Figure 6B**). Anti-S-RBD IgG titers in the lungs were elevated 7-28 dpi with a higher trend observed at 28dpi in females than males (*p*=0.09; **Figure 6C**). In the trachea, but not in nasal turbinate or lung homogenates, females had significantly greater anti-S-RBD IgG titers than males (*p*<0.05; **Figure 6D**). In summary, these data demonstrate that female hamsters develop greater systemic and local antiviral antibody responses compared with male hamsters during SARS-CoV-2 infection.

**Figure 6:**
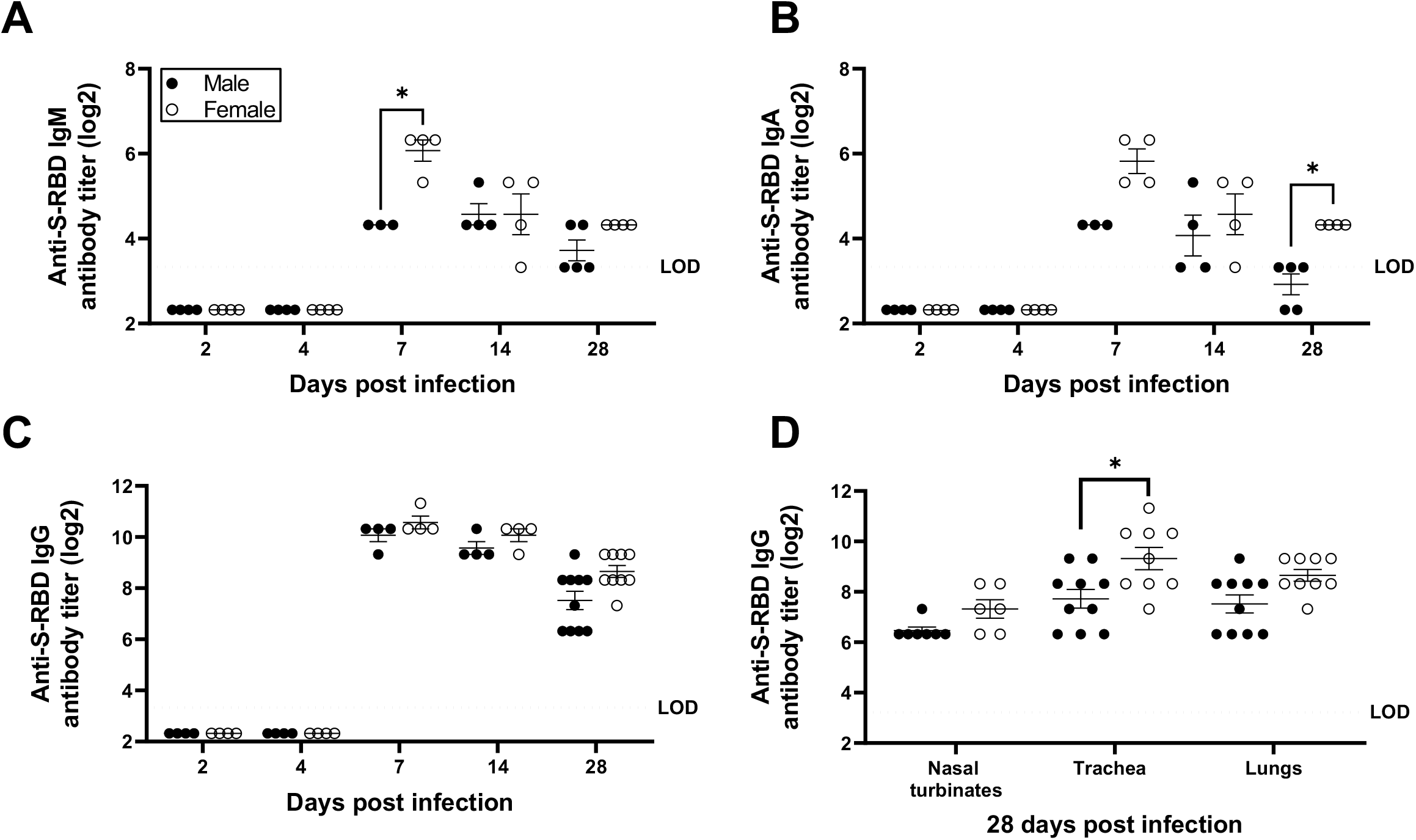
Antibody responses in the respiratory system of SARS-CoV-2 infected female hamsters were greater than males. Lung homogenates were prepared at different dpi and S-RBD-specific IgM (A), IgA (B), and IgG (C) antibodies were determined. Likewise, S-RBD-specific IgG antibodies were tested in the homogenates of nasal turbinates, trachea, and lungs at 28 dpi (D). Data represent mean ± standard error of the mean from one or two independent experiment(s) (n = 3-10/group) and significant differences between groups are denoted by asterisks (*p<0.05) based on two-way ANOVA (mixed-effects analysis) followed by Bonferroni’s multiple comparison test.

## Discussion

Sex differences in COVID-19 outcomes are well documented (9, 13). There is a critical need to develop accurate animal models that reflect the male-bias in disease outcomes to better understand the underlying mechanisms. We show that male hamsters suffer more systemic (body mass loss) and local (pulmonary pathology) symptoms of SARS-CoV-2 infection than females. We tested several potential mechanisms that could mediate male-biased outcomes from infection, including: 1) lack of estrogenic protection, 2) greater virus replication, 3) exacerbated cytokine responses, and 4) reduced humoral immunity. Our data reveal that females produce greater antibody responses, both locally in the respiratory tract as well as systemically in plasma, but if this causes female hamsters to suffer less severe outcomes from SARS-CoV-2 infection remains to be determined.

Clinical manifestations of SARS-CoV-2 infection in hamsters are typically mild, with reduced body mass after infection consistently observed (21, 24-26). Previous studies have shown that hamsters lose body mass after infection, reaching peak loss at 5 to 7 dpi, followed by recovery (24-26). Body mass loss in hamsters, regardless of age, has been associated with the dose of virus inoculum, with higher dose resulting in greater body mass loss (26, 41). Body mass loss also is influenced by age; older hamsters (i.e., 7 to 9 months old) had greater mass loss than younger animals (i.e., 4-6 weeks old) (21, 26). Sex is another factor impacting body mass loss, as a reliable clinical sign of disease in hamsters following SARS-CoV-2 infection. As reported in humans, older age and male sex are clinical variables associated with greater clinical manifestations of disease in hamsters.

A novel determinant of clinical disease that was utilized in the current study was unbiased, quantitative chest CT-imaging analysis. Previous reports describe chest CT findings in female SARS-CoV-2 infected hamsters only and show lung abnormalities, including multilobular ground-glass opacities (GGO) and consolidation (26), as observed in patients with COVID-19 (42). In the current study, CT-imaging revealed that multilobular GGO and consolidations were observed to a greater extent in male than female SARS-CoV-2-infected hamsters at 7 dpi. Whether the sexes differ in the recovery of pulmonary damage following infection requires greater consideration. There are a number of registered clinical trials of therapeutic E2 administration (NCT04359329 and NCT04539626) in COVID-19, which raised the question as to whether disease outcomes in male hamsters could be improved through administration of E2. In this study, pre-treatment of male hamsters with E2 prior to SARS-CoV-2 infection did not reduce either weight loss, observed histological damage to lung tissue or the observed multilobular GGO and consolidations.

SARS-CoV-2 replicates in the nasal turbinates, trachea, and lungs of infected golden Syrian hamsters (24, 25). Virus replication peaks in respiratory tissue within 2-4 dpi, with virus clearance typically occurring within one week (21, 24, 25). Viral RNA, however, is present in the lungs of infected hamsters beyond 7 dpi (21, 22, 25). We observed peak infectious virus load in nasal turbinates, trachea, and lungs at 2 dpi, with clearance by 7 dpi. After infectious virus had been cleared, viral RNA still remained detectable in the lungs up to 14 dpi. Previous studies have reported that while aged hamsters experience worse disease outcomes than young hamsters, virus titers in respiratory tissues are similar (21, 26). We further show that although young adult male hamsters experience worse disease outcomes than female hamsters, sex differences in virus titers in respiratory tissues are not observed.

During SARS-CoV-2 infection of hamsters, cytokine gene expression, including *Tnfα, Ifnα*, and *Ifnγ* in the nasal turbinates and lungs, is triggered at 2 dpi, peaks at 4 dpi, and returns to baseline by 7 dpi, but comparisons between males and females have not performed (24, 41, 43). Analyses of protein concentrations of cytokines in lung and spleen homogenates revealed no differences between males and females during the first week of infection. Although sex differences were not observed, concentrations of TNFα and IFNβ were positively and negatively, respectively, associated with virus replication in lungs, regardless of sex. Our findings suggest that cytokine production, either locally in the lungs or systemically in the spleen does not underlie sex differences in clinical manifestations of disease in hamsters and adds to the growing list of questions about the role of cytokines in the pathogenesis of SARS-CoV-2 in human populations. The possibility of differences in cellular infiltration into pulmonary tissue requires greater consideration, which will be feasible only when better reagents, including antibodies, become available for hamsters.

Studies have reported that both IgG and virus neutralizing antibodies are detected in serum from SARS-CoV-2-infected golden Syrian hamsters as early as 7 dpi and persist through 43 dpi (24, 35, 44). In the present study, females developed greater IgG responses against both SARS-CoV-2 wild type and variant S-RBD as well as antiviral nAb titers in both plasma and respiratory tissue homogenates than males. We also showed that mucosal IgA titers are greater in the lungs of female than male hamsters and are detectable as early as 7 dpi. Passive transfer of convalescent sera from infected to naïve hamsters as well as reinfection of previously infected hamsters carrying high antibody titers, have both been shown to provide protection by reducing virus titers in the respiratory tissues (24, 26). Likewise, hamster models of SARS-CoV-2 immunization have shown an inverse correlation between antibody responses and either virus titers in the respiratory tissues or body mass loss (45). These studies highlight the possible protective role of antibodies during SARS-CoV-2 infection, which may contribute to faster recovery in female than male hamsters.

Golden Syrian hamsters have already been successfully used in SARS-CoV-2 transmission studies (24, 25, 46), to compare routes of SARS-CoV-2 infection (41, 47), to evaluate convalescent plasma and monoclonal antibody therapy (24, 26, 48-50), and to test therapeutics and vaccines (23, 45). This model provides a unique opportunity to understand the kinetics of SARS-CoV-2 immunopathology not only systemically but also at the site of infection, the respiratory system. Sex as a biological variable should be considered in all studies utilizing golden Syrian hamsters for prophylactic and therapeutic treatments against SARS-CoV-2.

## Materials and Methods

### Viruses, cells, and viral proteins

Vero-E6-TMPRSS2 cells were cultured in complete cell growth medium (CM) comprising Dulbecco’s Modified Eagle Medium (DMEM) supplemented with 10% fetal bovine serum, 1mM glutamine, 1mM sodium pyruvate, and penicillin (100 U/mL) and streptomycin (100 μg/mL) antibiotics (51). The SARS-CoV-2 strain (SARS-C0V-2/USA-WA1/2020) was obtained from Biodefense and Emerging Infections Research Resources Repository (NR#52281, BEI Resources, VA, USA). SARS-CoV-2 stocks were generated by infecting VeroTMPRSS2 cells at a multiplicity of infection (MOI) of 0.01 TCID50s per cell and the infected cell culture supernatant was collected at 72 hours post infection clarified by centrifugation at 400 g for 10 minutes and then stored at -70C (51). SARS-CoV-2 recombinant spike receptor-binding domain (S-RBD) protein used for enzyme-linked immunosorbent assay (ELISA) was expressed and purified using methods described previously (14) or purchased from SinoBiologicals. To obtain whole inactivated SARS-CoV-2, VeroTMPRSS2 cells were infected at a MOI of 0.01 and the infected cell culture supernatant was collected at 72 hours post infection. Virus was inactivated by the addition of 0.05% beta-propiolactone (51) followed by incubation at 4C for 18 hours. The beta-propiolactone was inactivated by incubation at 37C for 2 hours and the inactivated virions were pelleted by ultracentrifugation at 25000*g* for 1h at 4^0^C and protein concentration was determined by BCA assay (Thermo Fisher Scientific).

### Animal experiments

Male and female golden Syrian hamsters (7-8 weeks of age) were purchased from Envigo (Haslett, MI). Animals were housed under standard housing conditions (68-76°F, 30-70% relative humidity, 12-12 light-dark cycle) in PNC cages (Allentown, NJ) with paper bedding (Teklad 7099 TEK-Fresh, Envigo, Indianapolis, IN) in an animal biological safety level 3 (ABSL-3) facility at the Johns Hopkins University-Koch Cancer Research Building. Animals were given nesting material (Enviropak, Lab Supply, Fort Worth, TX) and ad libitum RO water and feed (2018 SX Teklad, Envigo, Madison, WI). After 1-2 weeks of acclimation, animals (8-10 weeks of age) were inoculated with 10^5^ TCID_50_ (50% tissue culture infectious dose) of SARS-CoV-2 USA-WA1/2020) in 100μL DMEM (50μl/naris) through intranasal route under ketamine (60-80mg/kg) and xylazine (4-5mg/kg) anesthesia administered intraperitoneally. Control animals received equivalent volume of DMEM. Animals were randomly assigned to be euthanized at 2, 4, 7, 14, or 28-days post infection (dpi). Body mass was measured at the day of inoculation (baseline) and endpoint, with daily measurements up to 10 dpi and on 14, 21, and 28 dpi, when applicable per group. Blood samples were collected pre-inoculation (baseline) and at days 7, 14, 21, and 28 dpi, when applicable per group. Survival blood collection was performed on the sublingual vein, whereas terminal bleeding was done by cardiac puncture under isoflurane (500μl drop jar; Fluriso™, VetOne^®^, Boise, ID) anesthesia. Blood was collected into EDTA (survival and terminal) and/or sodium citrate tubes (terminal). Plasma was separated by blood centrifugation at 3500rpm, 15min at 4^0^C. After cardiac puncture, animals were humanely euthanized using a euthanasia solution (Euthasol^®^, Virbac, Fort Worth, TX). Nasal turbinates, trachea, and lung samples for antibody/cytokine assays and virus titration were snap frozen in liquid nitrogen and stored at -80^0^C.

### Determination of infectious virus titers and viral genome copies in tissue homogenates

To obtain tissue homogenates, DMEM with 100unit/mL penicillin and 100 μg/mL streptomycin was added (10% w/v) to tubes containing hamster nasal turbinate, lungs, and tracheal tissue samples. Lysing Matrix D beads were added to each tube and the samples were homogenized in a FastPrep-24 bench top bead beating system (MPBio) for 40sec at 6.0m/s, followed by centrifugation for 5min at 10,000g at room temperature. Samples were returned to ice and the supernatant was distributed equally into 2 tubes. To inactivate SARS-CoV-2, TritonX100 was added to one of the tubes to a final concentration of 0.5% and incubated at room temperature for 30 minutes. The homogenates were stored at -70^0^C.

Infectious virus titers in respiratory tissue homogenates were determined by TCID_50_ assay (14, 51). Briefly, tissue homogenates were 10-fold serially diluted in infection media (CM with 2.5% instead of 10% FBS), transferred in sextuplicate into the 96-well plates confluent with Vero-E6-TMPRSS2 cells, incubated at 37^0^C for 4 days, and stained with naphthol blue-black solution for visualization. The infectious virus titers in TCID_50_/mL were determined by Reed and Muench method. For detection of SARS-CoV-2 genome copies, RNA was extracted from lungs using the Qiagen viral RNA extraction kit (Qiagen) and reverse transcriptase PCR (qPCR) was performed as described (52).

### Computed tomography (CT) and image analysis

Live animals were imaged inside in-house developed; sealed biocontainment devices compliant with BSL-3, as previously reported (53). Seven days post-infection, SARS-CoV-2-infected males (n=13), females (n=14) and E2 treated (n=13) hamsters underwent chest CT using the nanoScan PET/CT (Mediso USA, MA, USA) small animal imager. CT images were visualized and analyzed using VivoQuant 2020 lung segmentation tool (Invicro, MA, USA) (54). Briefly, an entire lung volume (LV) was created, and volumes of interests (VOIs) were shaped around the pulmonary lesions using global thresholding for Hounsfield Units (HU) ≥ 0 and disease severity (CT score) was quantified as the percentage of diseased lung in each animal. The investigators were blinded to the group assignments.

### Hormone replacement and quantification

Estradiol (E2) capsules were prepared of Silastic Brand medical grade tubing (0.062 in. i.d. x 0.125 in. o.d.), 10 mm in length, sealed with Factor II 6382 RTV Silicone and Elastomer, and filled 5 mm with 17β-estradiol (55). Capsules were incubated overnight in sterile saline at 37°C prior to implantation. The E2 dosage was chosen because this size capsule has previously been shown to produce blood levels within the physiological range of E2 measured in intact female hamsters during early proestrus (when E2 levels are at their peak) (56, 57). Circulating concentrations of E2 were measured by a rodent estradiol ELISA kit as per manufacturer’s instructions (Calbiotech, CA).

### Antibody ELISAs

Hamster antibody ELISA protocol was modified from human COVID-19 antibody ELISA protocol described previously (14). ELISA plates (96-well plates, Immunol 4HBX, Thermo Fisher Scientific) were coated with either spike receptor binding domain (S-RBD) or whole inactivated SARS-CoV-2 proteins (2 μg/mL, 50μl/well) in 1X PBS and incubated at 4^0^C overnight. Coated plates were washed thrice with wash buffer (1X PBS + 0.1% Tween-20), blocked with 3% nonfat milk solution in wash buffer and incubated at room temperature for 1 hour. After incubation, blocking buffer was discarded and two-fold serially diluted plasma (starting at 1:100 dilution) or tissue homogenates (starting at 1:10 dilution) were added and plates were incubated at room temperature for 2 hours. After washing plates 3 times, HRP-conjugated secondary IgG (1:10000, Abcam, MA, USA), IgA (1:250, Brookwood Biomedical, AL, USA) or IgM (1:250, Brookwood Biomedical, AL, USA) antibodies were added. After addition of secondary IgG antibody plates were incubated in room temperature for 1 hour while for IgA and IgM antibodies, plates were incubated at 4^0^C overnight. Sample and antibody dilution were done in 1% nonfat milk solution in wash buffer. Following washing, reactions were developed by adding 100μl/well of SIGMAFAST OPD (o-phenylenediamine dihydrochloride) (MilliporeSigma) solution for 10 min, stopped using 3M hydrochloric acid (HCL) solution and plates were read at 490nm wavelength using ELISA plate reader (BioTek Instruments). The endpoint antibody titer was determined by using a cut-off value which is three-times the absorbance of first dilution of mock (uninfected) animal samples.

### Microneutralization assay

Heat inactivated (56^0^C, 35min) plasma samples were two-fold serially diluted in infection media (starting at 1:20 dilution) and incubated with 100 TCID_50_ of SARS-CoV-2. After 1-hour incubation at room temperature, plasma-virus mix was transferred into 96-well plate confluent with Vero-E6-TMPRSS2 cells in sextuplet. After 6 hours, inocula were removed, fresh infection media was added, and plates were incubated at 37^0^C for 2 days. Cells were fixed with 4% formaldehyde, stained with Napthol blue black solution and neutralizing antibody titer was calculated as described (14).

### Cytokine ELISAs

Cytokine concentrations in TritonX100 inactivated lung and spleen homogenates were determined by individual ELISA kits for hamster IFN-α (mybiosource.com; MBS010919) IFN-β (mybiosource.com; MBS014227), TNF-α (mybiosource.com; MBS046042), IL-1β (mybiosource.com; MBS283040), IFNγ (ARP; EHA0005), IL-10 (ARP; EHA0008), and IL-6 (ARP; EHA0006) as per the manufacturer’s instructions. Samples were pre-diluted 1:5 to 1:10 as necessary in the appropriate kit sample dilution buffer. Total protein in the homogenates were measured by BCA assay (Thermo Fisher Scientific).

### Statistical Analyses

Statistical analyses were done in GraphPad Prism 9. Changes in body mass were compared using two-way repeated measures ANOVA followed by Bonferroni’s multiple comparison test. Chest CT scores were compared by unpaired Mann-Whitney test. E2 concentration were compared by two-tailed unpaired t-test. Virus titers and antibody responses were log transformed and compared using two-way ANOVA or mixed-effects analysis followed by Bonferroni’s multiple comparison test. Cytokine concentrations were normalized to total protein content in lung homogenates and compared using two-Way ANOVA. Associations between cytokines and virus titers in lungs were conducted using Spearman correlational analyses. Differences were considered to be significant at *p*<0.05.

## Data availability

All data will be made publicly available upon publication and upon request for peer review.

## Acknowledgements

We are grateful to the Johns Hopkins School of Medicine Vice Dean of Research, Dr. Antony Rosen, for providing research funds to develop this model and conduct this research. We also thank the Johns Hopkins COVID-19 Hamster Study Group members for weekly discussions and participation in these studies, particularly the veterinarians and animal care staff who ensured proper care of all animals in this study. AP would like to dedicate this manuscript to the memory of R. Mark Buller, whose collaborations on the golden Syrian hamster model for SARS-CoV infection formed the basis for this study.

## Funding

These studies were supported through the generosity of the collective community of donors to the Johns Hopkins University School of Medicine for COVID research with supplemental funds from The Johns Hopkins Center of Excellence in Influenza Research and Surveillance (CEIRS; HHSN272201400007C; AP, SLK), the NIH/NCI COVID-19 Serology Center of Excellence U54CA260492 (SLK), the NIH/ORWH/NIA Specialized Center of Research Excellence in Sex Differences U54AG062333 (SLK), R01AI153349 (SKJ), support from the Center for Infection and Inflammation Imaging Research (Johns Hopkins University), and NIH T32OD011089 (JLM).

## Conflicts of interest

The authors report none.

## Contributions

P.C.K., J.L.M., A.P., S.K.J., and S.L.K. conceptualized and designed the study. S.D., C.A.R-B., P.S.C., J.S.V., A.A.O., K.F., A.L.M., F.M., M.W., F.R.D., K.A.M-P and S.K.J\ designed and performed animal experiments. C.A.R-B, K.F, F.M., and M.W performed chest CT scans. A.P., R.Z., P.M., A.J., N.M, and M.J.B. grew virus, did virus quantification, and produced antigens required to run ELISAs. S.D., K.L., P.S.C., A.L.M., and A.P. performed antibody assays. S.D., P.S.C., J.R.C., M.M.L., D.Q., and P.C.K. performed cytokine assays. S.D. and C.A.R-B ran statistical analyses on data. The Study Group Members performed animal experiments, tissue processing, and data management. S.D. and S.L.K. wrote the manuscript with input from all authors. All authors read and provided edits to drafts and approved the final submission.

## Figure legends

**Supplementary Figure 1:** Representative transverse chest CT of five females, placebo-treated males, and E2-treated male hamsters at 7 dpi. Multiple bilateral and peripheric ground-glass opacities (GGO) and mixed GGO with consolidations are the hallmarks findings at the peak of lung disease.

**Supplementary Figure 2:** Kinetics of cytokine concentrations (pg/mg total protein)in the lungs of SARS-CoV-2 infected hamsters. Male and female golden Syrian hamsters were infected with 10^5^ TCID_50_ of SARS-CoV-2. Subsets of animals were euthanized at different dpi and IL-1β (A), TNF-α (B), IL-6 (C), IFN-α (D), IFN-β (E), and IFN-γ (F) concentrations were determined in the lungs by ELISA. Mock-infected animal samples from 2-, 4-, or 7-days post infection (dpi) were not statistically different and were combined and presented together as 0 dpi. Data represent mean ± standard error of the mean from one or two independent experiments (n = 6-12/group) with significant differences between groups denoted by asterisks (*p<0.05) based on one-way ANOVA followed by Dunnett’s multiple comparisons test.

**Supplementary Figure 3:** Associations between concentrations (pg/mg total protein) of IL-1β (A), TNF-α (B), IL-6 (C), IFN-α (D), IFN-β (E), and IFN-γ (F) and virus titers in lungs collected 2 days post infection (dpi). Data were analyzed with Spearman correlation analyses with significant associations represented with the R statistic and associated p-value.

**Supplementary Figure 4:** Associations between concentrations (pg/mg total protein) of IL-1β (A), TNF-α (B), IL-6 (C), IFN-α (D), IFN-β (E), and IFN-γ (F) and virus titers in lungs collected 4 days post infection (dpi). Data were analyzed with Spearman correlation analyses with significant associations represented with the R statistic and associated p-value.

**Supplementary Table 1:** Concentrations (pg/mg total protein) of cytokines in the lungs and spleen of male and female hamsters at different days post infection (dpi). Mock-infected animal samples from different dpi were pooled and used as 0 dpi. Data are presented as the mean ± standard error of the mean from one or two independent experiments (n = 6-12/group) with no significant differences observed between the groups based on two-way ANOVA (mixed-effects analysis) followed by Bonferroni’s multiple comparison test.

